# DNMT3A-NPM1 mutated acute myeloid leukaemia shows sensitivity to a PARP1 inhibitor combined with daunorubicin in an in vitro model

**DOI:** 10.1101/362103

**Authors:** Grigore Gafencu, Valentina Pileczki, Ancuta Jurj, Lorand Magdo, Cristina Selicean, Roxana Ola, Gabriel Ghiaur, Ioana Berindan-Neagoe, Ciprian Tomuleasa

## Abstract

Acute myeloid leukaemia is a neoplasia in need of new treatment approaches. PARP inhibitors are a class of targeted therapeutics for cancer that disrupts dysfunctional DNA damage response in various neoplasia. MLL-AF9 mutated leukaemias are sensitive to combinations of PARP inhibitors and cytotoxic drugs. Moreover, DNMT3A and NPM1 mutations are linked to dysfunctions in DNA damage response. Therefore, we investigated if DNMT3A-NPM1 mutated AML cell line is sensible to PARP inhibitors combined with anthracyclines. Our results show that DNMT3A-NPM1 mutated AML is as sensible to combinations of PARP inhibitors and anthracyclines as MLL-AF9 mutated leukaemias, in an in vitro setting.

---

Acute myeloid leukaemia (AML) is a malignant disorder characterised by the accumulation of genetic aberrations in the myeloid haematopoiesis linage, that can be risk classified according to specific gene mutations (1). Understanding the AML genome promises new therapeutic strategies that involve synthetic lethal interactions (2). Thus, in this study we compare the effects of combining olaparib, a poly (ADP-ribose) polymerase (PARP) inhibitor that induces synthetic lethality in malignancies bearing homologous recombination deficiencies (2) and daunorubicin (ODC) with the standard daunorubicin-cytarabine (CDR) that models the “7+3” induction regimens. Two AML cell lines were chosen: OCI/AML3 and THP-1. OCI/AML3 is characterised by a common and impactful co-occurrence of mutations in both nucleophosmin gene (*NPM1*), involved in DNA single strand break repairs and DNA methyltransferase 3 alpha gene (*DNMT3A*), involved in resistance to anthracycline-induced DNA damage (3)(4)(5). THP-1 is characterised by mutations and deletions in *PTEN*, *MLL*-*AF9*, *MLLT3*, *TP73*, *CDKN2A/B* (6), which makes it susceptible to the effects of olaparib, particularly through the presence of a partial deletion in *PTEN* gene (2).

Firstly, we subjected OCI/AML3 (DSMZ, Braunschweig, Germany) and THP1 (ATCC, Manassas, USA) cells to an in vitro drug treatment, by plating them in 96-well plates, at 10^4^cells/200μL/well in 2 types of media: 80% alpha-MEM (Invitrogen, Paisley, UK) with 20% FBS (Invitrogen, Paisley, UK) for OCI/AML3 and RPMI1640 (Invitrogen, Paisley, UK) with 10% FBS, 2 mM L-glutamine for THP1 for 48h/72h with 37.5μM olaparib (Selleckchem, Houston, USA), 100μM cytarabine (Sigma Aldrich, Taufkirchen, Germany), 1.4μM daunorubicin (Sigma Aldrich, Taufkirchen, Germany) alone or in combination. Cells were maintained in cell culture incubators at 37°C and 5%CO_2_ atmosphere. Cell proliferation was assessed with the CellTiter 96^®^ AQueous Non-Radioactive Cell Proliferation Assay (Promega, Madison, USA) and analysed with a BioTek Synergy H1 Hybrid Multi-Mode Reader (BioTek Instruments, Winooski, USA). Cell cycle arrest was assessed using a flow cytometry-based approach at 48h of treatment, as described elsewhere (7). DNA double breaks (DSB) levels after 48h and 72h treatments were determined as an increase in phosphoSer139 *γ*H2AX foci, a DSB marker, using flow cytometry method as described elsewhere (8). Gene expression analysis for DNA damage associated genes, *ATM*, *RAD51*, *LIG3*, *LIG4*, *PARP1*, *PTEN* and *B2M*, as internal normaliser, was performed using qRT-PCR as described elsewhere (9) and custom-made primers. All the experiments were conducted in triplicate and represented as means ±SEM/SD. We analysed all the data using the GraphPad Prism 6 software suite and applied ANOVA statistics and Dunnett or Sydak tests. Results were considered significant for p values ≤ 0.05.

Moving to results, ODC was as effective as CDR and daunorubicin in arresting the relative viability of the treated cells (Fig. 1A – OCI/AML3: 54.9% *vs*. 56.1% vs. 58.13%; THP1:38.5% *vs*. 30.1% vs. 31.57%, Dunnett test for multiple comparisons, *p*>0.982). Moreover, ODC reproduced the effects of CDR and daunorubicin monotherapy on the distribution of cells in each cell cycle stage, regardless of cell line tested (Fig.1B Dunnett’s multiple comparisons test *p*>0.05). Furthermore, at 48h time point, there was no significant difference in DSB induction by ODC, CDR or daunorubicin monotherapy, regardless of treated cell line. However, at the 72h time point, significant differences were observed between ODC, CDR and daunorubicin DSB production, regardless of cell line (Fig.1C relative signal intensity - OCI/AML3 - ODC vs CDR vs daunorubicin: 131% *vs*. 122% vs. 141.2%; THP1 - ODC vs CDR vs daunorubicin: 127% *vs*. 146% vs. 136.6%, Sidak’s test for multiple comparisons, *p*=0.0211).

**Figure 1.**
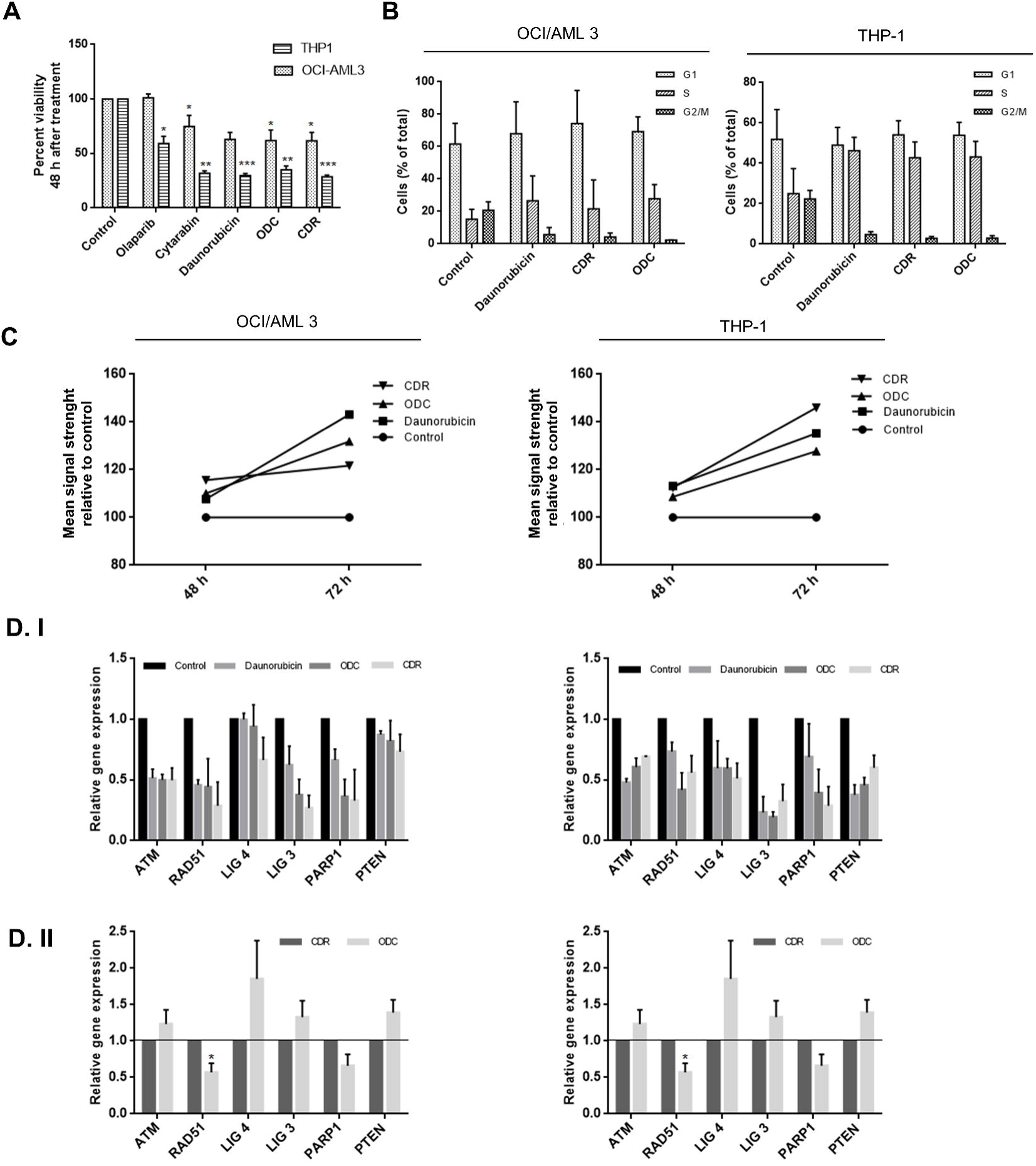
Effects of the individual and combined treatments of 37.5 μM olaparib, 1.4 μM daunorubicin and 100 μM cytarabine on OCI/AML3 and THP-1, at both functional and gene expression level. A) MTS cell viability analysis performed 48 h after the treatment. Data is represented as mean ± SD. ^∗^p<0.05, ^∗∗^p<0.001, ^∗∗∗^p<0.0001. B) Effects of ODC and CDR in both OCI/AML3 and THP-1 cell lines on G2/M cell cycle arrest after 48 h. Experiments were performed in triplicates and results depicted as mean ±SD. C) Time dependent DNA DSB accumulation. Mean signal strength generated by the antibody-Alexa Fluor 488 labeled phosphoSer139 *γ*H2AX foci in treated and untreated cells, relative to untreated control. Data from 48 and 72h time points. D. I) qRT-PCR gene expression profile of key components in PARP signaling pathway related to the previous established treatment strategy, 48h drug treatment. All the treatments were reported to the untreated control as a baseline. Three biological replicates were analyzed and expressed as mean ± SEM. ^∗^p<0.001 D. II) qRT-PCR gene expression profile of key components in PARP signaling pathway related to the previous established treatment strategy, 48h drug treatment. CDR was used as base line for the comparisons. Three biological replicates were analyzed and expressed as mean ± SEM. ^∗^p<0.001

ODC and CDR treatments induced downregulations in all the evaluated genes, when compared to the untreated control, in both cell lines. For OCI/AML3, daunorubicin monotherapy did not lower expression levels more than ODC and CDR for any studied genes. However, for THP1, daunorubicin alone downregulated *LIG3* and *PTEN* more than ODC and CDR (Fig.1 D. I). When comparing the gene expression for OCI/AML3 cells treated with ODC relatively to the ones treated with CDR, *ATM*, *PTEN*, *LIG3*, *LIG4* where found to be upregulated, and *RAD51* and *PARP1* downregulated. When the same analysis was performed for THP-1 cells, all the genes registered as downregulated (Fig 1 D. II). Gene expression levels induced by ODC and CDR were significantly different from those generated by daunorubicin monotherapy, regardless of cell line (Dunnett’s multiple comparisons test, p <0.05).

To conclude, the effects produced by ODC on blast proliferation, cell cycle and DNA damage levels mirrored the ones induced by CDR and daunorubicin regardless of the cell lines tested, but ODC generated gene expression patterns different from daunorubicin monotherapy and CDR. Our results show that NPM1-DNMT3A mutated AML is susceptible to the action of ODC in this in vitro setting and reinforce the reports that so does THP1, too (10). This phenomenon can be attributed to increased amounts of DSB, but also probably, to the effects of ODC on the expression of *RAD51* and *PARP1* genes on both cell lines. This observation is intriguing since inhibition of PARP1 delays the onset of starvation and ROS-induced autophagy (11) with potential effects on blast survival. The in vitro effects of ODC on these AML cell lines suggest that combining PARP inhibitors with anthracyclines could capitalise on two defective signalling machineries in AML: DNA repair and probably, autophagy. This is of potential clinical impact, as it can ease the side effect burden of AML treatment by substituting cytarabine with olaparib in treating patients with *NPM1-DNMT3A* mutated AML without jeopardising efficacy.

## Acknowledgements

This paper was funded by: “Iuliu Hatieganu” University of Medicine and Pharmacy internal student grant n°4995/10/08.03.2016 awarded to Grigore Gafencu and 2 Romanian Government research grants awarded to Ciprian Tomuleasa (PNII-RU-TE-2014-4-1783) and Roxana Ola (PN-II-RU-TE-2014-4-2951). Grigore Gafencu and Valentina Pileczki contributed equally to the current paper; Grigore Gafencu, Valentina Pileczki, Ancuţa Jurj, Lorand Magdo performed the experiments; Grigore Gafencu, Valentina Pileczki, Cristina Selicean, Ciprian Tomuleasa, Gabriel Ghiaur, Roxana Ola designed the experiments; Grigore Gafencu, Valentina Pileczki, Ciprian Tomuleasa, Gabriel Ghiaur, Ioana Berindan-Neagoe wrote the manuscript; all the authors were involved in the final approval of manuscript.

